# eXclusionarY: Ten years later, where are the sex chromosomes in GWAS?

**DOI:** 10.1101/2023.02.03.526992

**Authors:** Lei Sun, Zhong Wang, Tianyuan Lu, Teri A. Manolio, Andrew D. Paterson

## Abstract

Ten years ago, a detailed analysis of genome-wide association studies showed that only 33% of the studies included the X chromosome. Multiple recommendations were made to combat eXclusion. Here we re-surveyed the research landscape to determine if these earlier recommendations had been translated. Unfortunately, among the summary statistics reported in 2021 in the NHGRI-EBI GWAS catalog, only 25% provided results for the X chromosome and 3% for the Y chromosome, suggesting that the eXclusion phenomenon documented earlier not only persists but has also expanded into an eXclusionarY problem. Normalizing by physical length of the chromosome, the average number of studies published until 11/29/22 with genome-wide significant findings on the X chromosome is ~1 study/Mb. In contrast, it ranges from ~6 to ~16 studies/Mb for chromosomes 4 and 19, respectively. Compared with the autosomal growth rate of ~0.086 studies/Mb/year over the last decade, studies of the X chromosome grew at less than one-seventh that rate, only ~0.012 studies/Mb/year. Among the studies that reported significant association on the X chromosome, there were extreme heterogeneities in how they analyzed the data and documented the results, suggesting the need for guidelines. Not surprisingly, among the 430 scores sampled from the PolyGenic Score catalog, 0% contained weights for sex chromosomal SNPs. To overcome the dearth of sex chromosome analyses, we provide five sets of recommendations and future directions. Finally, until the sex chromosomes are included in a whole-genome study, instead of GWAS, we propose they be more properly referred to as “AWAS” for “autosome-wide scan”.

---

In the ten years since Wise et al. (2013)^1^ brought the eXclusion of the X chromosome from genome-wide association studies (GWAS) to the attention of the community, little has improved regarding the analysis and reporting of the sex chromosomal variants in GWAS^2–4^. The X chromosome accounts for ~5% of the haploid genome and carries ~800 protein-coding genes. However, to date (November 2022), even after the call for including the X chromosome in GWAS by ^1^, approximately only 0.5% of associated SNPs in the NHGRI-EBI GWAS catalog^5,6^ are on the X chromosome, a 10-fold paucity compared to the autosomes.

The paucity of research on the sex chromosomes includes both the X and Y chromosomes. For the Y chromosome^7^, as of November 29, 2022, only nine out of 447,939 associations reported in NHGRI-EBI GWAS Catalog^5,6^ belong to the Y chromosome. Coverage is scarce on GWAS arrays for the male-only Y chromosome, in part due to repetitive sequences that makes variant calling difficult. If Y chromosomal variants are available in the non-pseudo-autosomal region (NPR), they can be analyzed using existing methods. However, there appears to be “a lack of will” to do so^8^.

For the X chromosome, there are multiple analytical challenges^9–14^, including i) a male has one copy of the X chromosome while a female has two, in contrast to the autosomes, ii) the X chromosome in male germ cells only recombines with the Y chromosome in the pseudo-autosomal regions (PARs) but not in the NPR, iii) in contrast to males, the two copies in female germ cells recombine across the entire X chromosome, iv) the two female copies are also subject to X-inactivation (i.e. X chromosome dosage compensation), v) the X-inactivation status at the population level can be random, skewed, or absent (i.e. X-inactivation escape), and vi) the true X-inactivation status at the individual level cannot be derived from GWAS data alone.

Thus, the existing bioinformatic, statistical and machine learning methods developed specifically for the autosomes are not suitable for the sex chromosomes. For example, most bioinformatic tools are autosome-centric, meaning that even if the sex chromosomes were included in the pipelines, tool developments were not tailored for the sex chromosomes^11^. These include variant calling^15,16^, data quality control (QC) prior to imputation^17,18^ (e.g. cryptic relatedness^19,20^, Hardy-Weinberg equilibrium (HWE)^21,22^), and imputation^15,23–30^. Similarly, most association methods are not tailored for the sex chromosomes, including population stratification via principal component analysis (PCA)^31^, and the association methodology itself^32–36^. Finally, the recent polygenic risk score (PRS)-based disease risk prediction methods^37–40^ rarely include the sex chromosomes. Even if the sex chromosomes were examined, the potential sex difference in, for example, linkage equilibrium, is rarely evaluated or acknowledged, even though earlier work has shown sex-specific linkage^41–44^.

Back in 2013, after examining 743 GWAS papers published between January 2010 and December 2011 and in the NHGRI GWAS Catalog^45^, Wise and colleagues noted that only ~33% GWAS included the X chromosome^1^; the Y chromosome was not explicitly examined, though it is implicitly involved in the X chromosome through the PARs. Additionally, the authors commented on QC and power concerns, including poorer coverage of the X chromosome in early GWAS arrays and lower genotyping and imputation accuracy as compared to autosomes, as well as X-inactivation-related analytical complexities that may reduce power of an association study. Finally, the authors concluded that “many interesting biological insights could be revealed if we end the exclusion of the X chromosome in future GWAS”.

Thus, ten years later, we first re-surveyed the research landscape to determine if the earlier recommendations of including the X (and Y) chromosomes in GWAS had been translated into changes in practice. Second, as genotyping and sequencing technologies have also evolved, including imputation panels based on next-generation sequencing data^15,46^, we then scanned the literature for emerging issues and insights. Finally, we make new recommendations.

## Sex Chromosome Results in the NHGRI-EBI GWAS and PolyGenic Score (PGS) Catalogs

### Lack of X and Y SNP-trait associations in the NHGRI-EBI GWAS catalog

As of Nov 29, 2022, the NHGRI-EBI GWAS Catalog^5,6^ contained 6,130 published studies, of which 4,208 reported at least one genome-wide significant association (p-value < 5 × 10^-8^)^47^. However, only 186 studies (4.4%) had signals on the X chromosome (Figure 1A). In contrast, chromosome 21 had twice the number of signals (418 studies; 9.9%), despite being less than one-third the length (Figure 1A).

**Figure 1.**
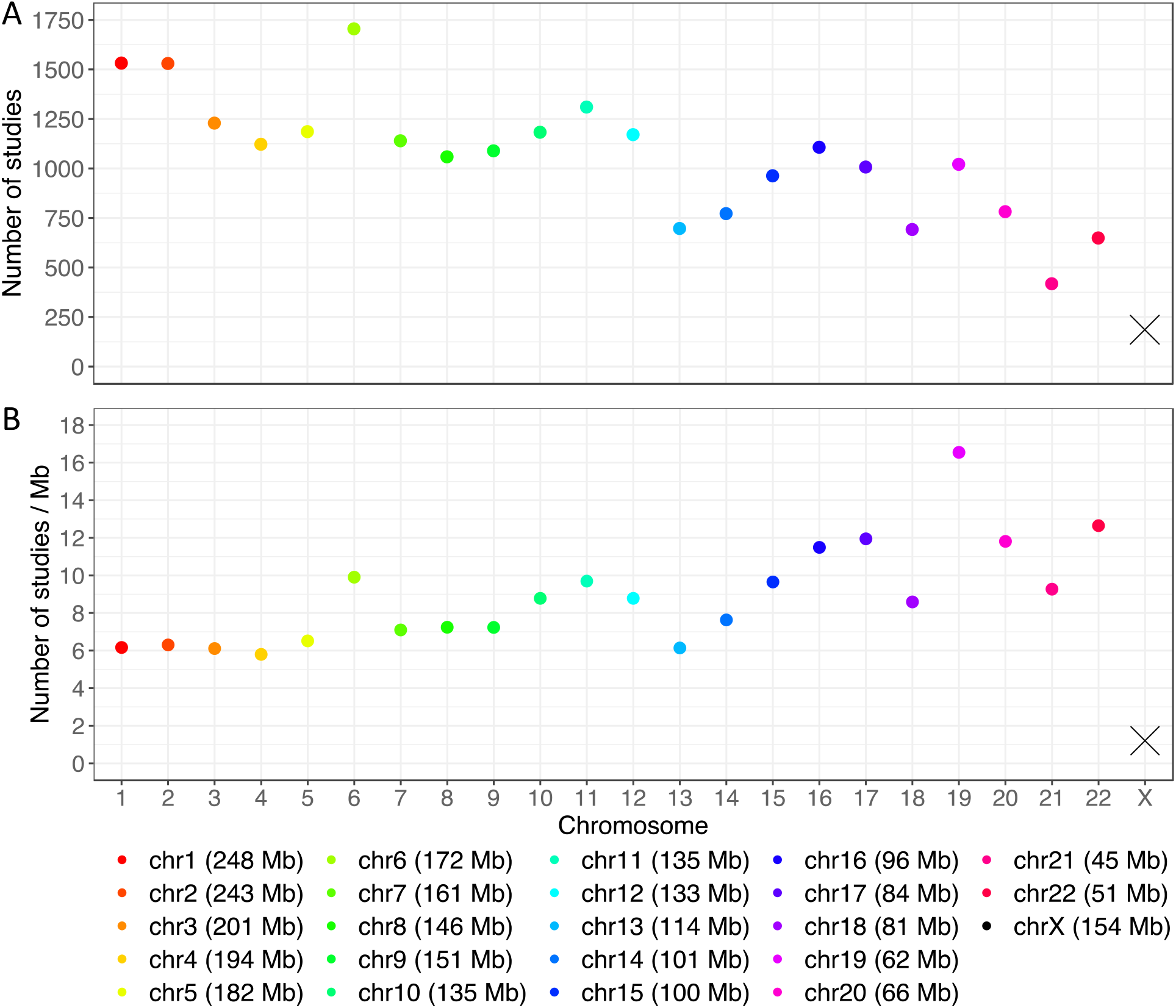
(A) Total number of studies and (B) average number of studies per Mb reporting at least one genome-wide significant finding (p-value < 5 × 10^-8^) stratified by chromosome, from the NHGRI-EBI GWAS Catalog up to November 29, 2022. Genetic associations were indexed by unique PubMed IDs. Studies reporting associations with multiple traits were only counted once.

Before investigating how often the X chromosome was analyzed to begin with (in the next section), we first normalized each chromosome by its physical length (Figure 1B). It is clear that signal densities vary across the autosomes. However, the most striking feature is the continued paucity of signals on the X chromosome since 2010-2011^1^.

To investigate whether the 2013 recommendation to include the X chromosome in GWAS had an impact on the practice of our field, we examined temporal changes. Figure 2 shows the average number of studies per Mb with at least one genome-wide significant finding, separately for the autosomes and the X chromosome, from prior to 2008 to November 29, 2022. Unfortunately, the gap between the autosomes and the X chromosome appears to be widening in recent years. Between 2009 and 2021, the average number of studies with genome-wide significant findings on the X chromosome grew at approximately 0.012 studies/Mb/year, remaining below 0.3/Mb every year (Figure 2 and Supplementary Figure S2). In contrast, the numbers increased consistently for the autosomes, by approximately 0.086 studies/Mb/year.

**Figure 2.**
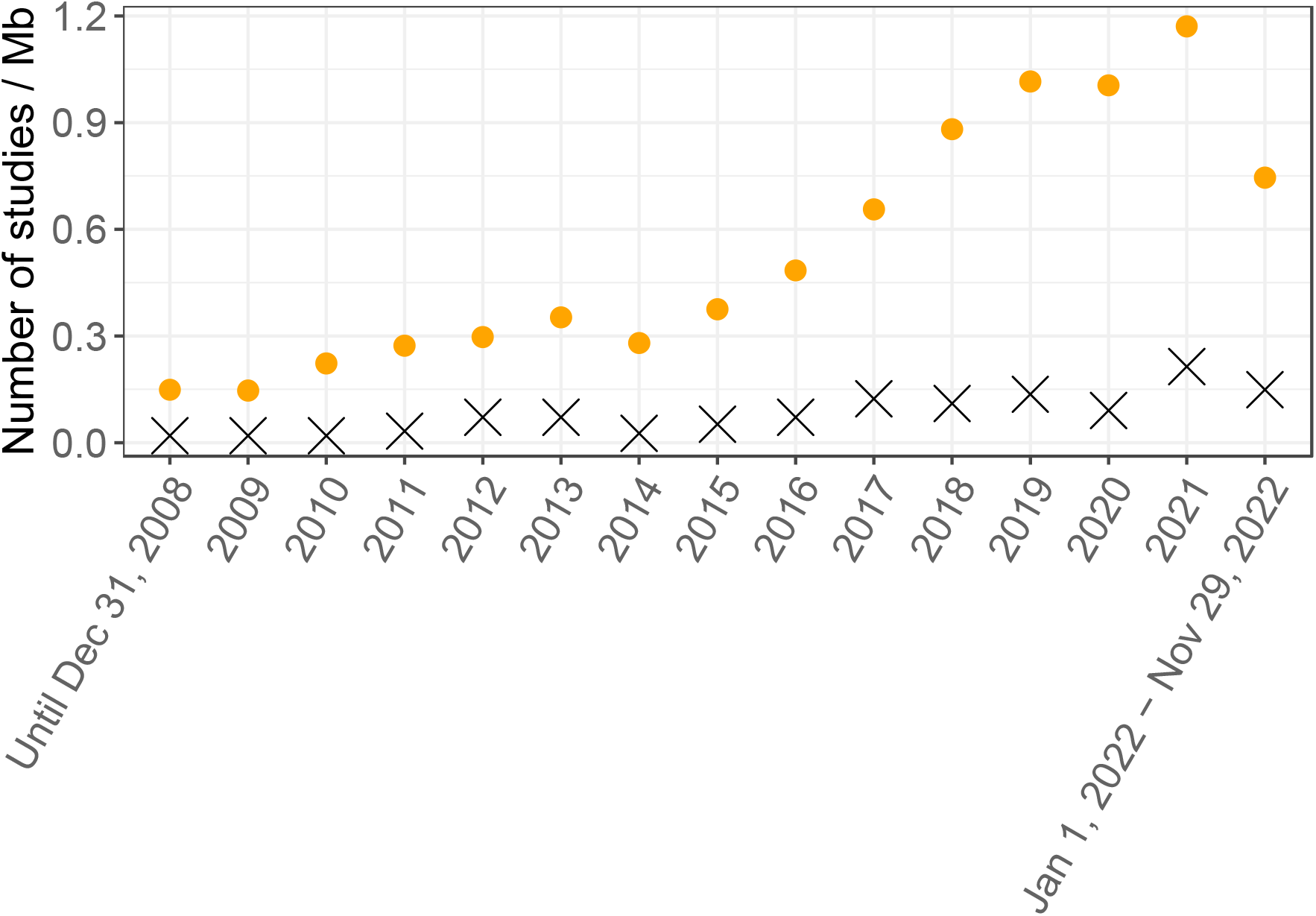
Average number of studies per Mb reporting at least one genome-wide significant finding (p-value < 5 × 10^-8^) over time, separately for the autosomes and X chromosome, from the NHGRI-EBI GWAS Catalog. Genetic associations were indexed by unique PubMed IDs. Studies reporting associations with multiple traits were only counted once.

Genetic associations on the Y chromosome were even more rarely documented. Out of all 447,939 associations (p-value < 1 × 10^-5^), only nine, arising from two studies, were on the Y chromosome; among the 293,170 genome-wide significant findings, only one was from the Y chromosome. These results indicate that the eXclusion phenomenon documented earlier not only persists but has also expanded into an eXclusionarY problem, where both sex chromosomes have been routinely neglected in whole-genome studies.

### Lack of X and Y chromosome results in GWAS summary statistics in the NHGRI-EBI GWAS catalog

To address whether lack of sex chromosomal GWAS results were due to lack of appropriate (or any) analysis of the sex chromosomes, we calculated the proportion of GWAS summary statistics that included sex chromosome results, regardless if there were significant findings.

There were 19,935 GWAS summary statistics published in 2021 and posted at the NHGRI-EBI GWAS catalog^5,6^ (Resources). These GWAS submissions came from 136 publications, of which most provided 1-2 sets of summary statistics, but four provided > 1,000 sets (Supplementary Data S1). To avoid analyzing multiple submissions from the same publication, we randomly selected one submission from each of the 136 publications (Supplementary Data S1).

Out the 136 GWAS summary statistics, only 34 (25%) contained X chromosome results (of the 34, only 4 also included Y chromosome results), which is less than the 33% based on the survey of GWAS conducted in 2010 and 2011^1^. Thus, eXclusion has become more rather than less prevalent, contrary to the intent of the initial commentary!

If we assume that the 136 studies with summary statistics in the NHGRI-EBI GWAS catalog are a random sample of all GWAS in 2021, and recall from the previous section that there is a six-fold difference in the average findings between chromosome 1 and the X chromosome (Figure 1B), it is then reasonable to hypothesize that much of the paucity would be resolved if the X chromosome were actually analyzed across all GWAS.

Variation in single nucleotide diversity^48,49^ can be a contributing factor. For example, chromosome 19 has the highest density of single nucleotide variations of 43.21/kb (based on the 1000 Genomes Project) among all chromosomes, while chromosome 1 has the lowest of all autosomes at 36.46/kb. However, cumulatively as of December 2022, there is no statistically significant linear relationship (slope = 0.54; p-value = 0.13; Figure 3) between nucleotide diversity and the average number of genome-wide significant findings among the autosomes. Even if we were willing to extrapolate the linearly fitted line to 30.16/kb, the nucleotide diversity of the X chromosome, the expected research yield on the X chromosome is 3.47/Mb, almost thrice the actual output of 1.21/Mb.

**Figure 3.**
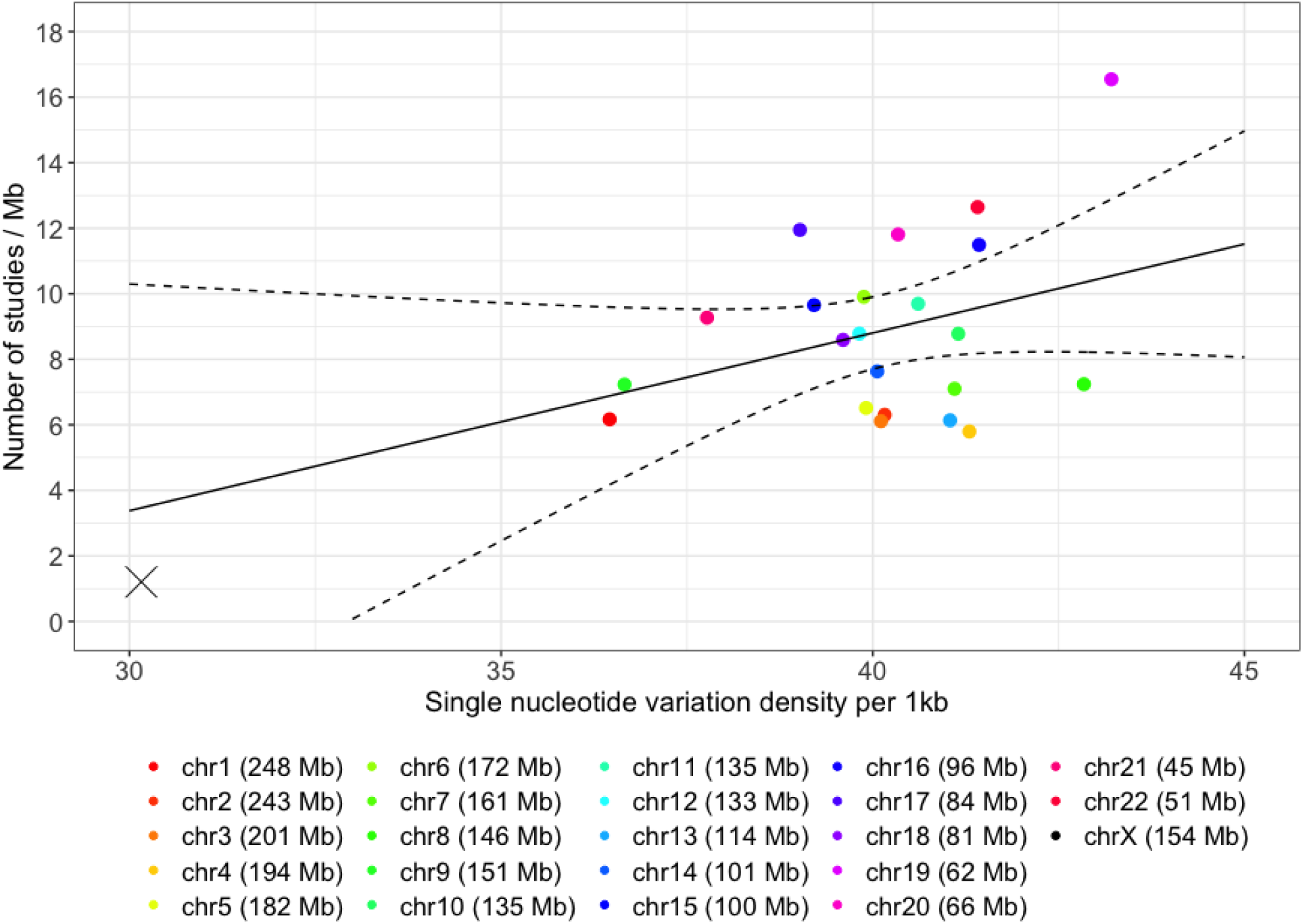
Average number of studies per Mb reporting at least one genome-wide significant finding (p-value < 5 × 10^-8^) per chromosome, from the NHGRI-EBI GWAS Catalog cumulatively up to November 29, 2022, compared to chromosome-specific nucleotide diversity^49^. Genetic associations were indexed by unique PubMed IDs. Studies reporting associations with multiple traits were only counted once. The solid slope was fitted using the autosomal data only, and the dashed curves are the 95% confidence bands.

Based on high-coverage whole-genome sequencing of TOPMed^15^ cohorts, the X chromosome has lower density of variants in coding sequences, as compared to the autosomes^50^. This can be an additional contributing factor to the paucity of signals on the X chromosome, but the sequenced individuals in TOPMed are typically middle-aged adults, which may result in selection bias against X chromosomal coding variants.

### Lack of X and Y chromosome results in the PolyGenic Score (PGS) catalog

As expected, there is also a lack of sex chromosome results in the PGS catalog. We downloaded PGS scoring files from the PGS catalog^51^ (Resources), focusing on the 430 files (PGS001802 to PGS002231) all uploaded on January 10, 2022. Unsurprisingly, none of the 430 files contained any results from the sex chromosomes, confirming the current eXclusionarY practice in PGS research as well.

## Other Emerging eXclusionarY Issues: Quality Control, Association Analysis and Reporting, Results Interpretation, the Y chromosome, and Clinical Implications

### Quality control

In addition to the QC discussed in ^1^, many data quality pipelines and imputation tools^15–18,23–30^ have been developed for GWAS. However, most are autosome-centric, ignoring the sex chromosomes either explicitly or implicitly. In 2014, ^11^ highlighted “the steps in which the X chromosome requires specific attention, and [gave] tentative advice for each of these”, including sex-stratified MAFs and missing rates, as well as testing for differential missingness. However, these recommendations have not been followed in practice. For example, sex-specific variant call rates are rarely reported.

### Sex difference in minor allele frequency (sdMAF) as QC revisited

Checking for sex difference in minor allele frequency is rarely formed as part of GWAS QC. However, it was already noted a decade ago that “MAF checks might need to be conducted separately for the X chromosome because the expected frequencies are sex dependent,” based on an informal poll of leading statistical geneticists working in GWAS in 2013^1^. Others also suggested to include a sdMAF test as part of the QC for the X chromosome^11^. However, a recent work has shown that there are possible causes of sdMAF: genotyping errors and biology^52^. Delineating the two causes for each X chromosomal SNP is not straightforward, creating challenges in QC pipelines.

The recent study analyzed the high-coverage whole genome sequencing data of the 1000 Genomes Project^49^ and gnomAD v3.1.2^53^, and identified many SNPs with genome-wide significant sdMAF across the X chromosome, particular at the boundaries between PAR and NPR^52^. Further, the study concluded that region-specific sdMAF at the PAR-NPR boundaries is likely a biological phenomenon, possibly due to sex-specific linkage^41,43,44^. This illustrates the challenges of including sdMAF as a QC measure.

As sdMAF is statistically equivalent to GWAS of sex, there is also a connection between the sdMAF study^52^ and a recent GWAS of sex^54^. This GWAS of sex used data from 2.46 million customers of 23andMe but did not examine the X chromosome^54^. Although their main conclusion was that sdMAF is a result of participation bias, they also noted that 55% of their significant findings on the autosomes are likely results of genotyping errors, further illustrating the importance and challenges of separating genotyping errors from biology (and other causes) that could lead to sdMAF.

### Hardy-Weinberg equilibrium (HWE) test as QC revisited

Departure from Hardy-Weinberg equilibrium is routinely used as part of GWAS QC for autosomes^17^, as SNPs with severe Hardy-Weinberg disequilibrium (HWD) are typically believed to have genotyping errors^55^. However, how to evaluate HWE for the X chromosome is unclear, and it remains debatable whether testing for HWD should be used at all as part of data QC, both of which we discuss next.

The standard HWE test is Pearson’s 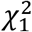 test, testing for the difference between the observed and expected genotype counts based on HWE^56,57^. This test is typically applied to sex-combined genotype counts, which is reasonable for an autosomal SNP. But applying such a HWE test to an X chromosomal PAR or NPR SNP requires additional considerations^58^. For example, ^11^ recommended performing the HWE test using only females.

Alternatively, ^21,59^ suggested using both females and males, and they proposed a new HWE test for a NPR SNP that includes the deviation of male genotype counts from the expected, based on sex-pooled allele frequency estimate. However, this alternative test has been shown to be simultaneously testing for HWD in females, and sdMAF between males and females^58^. Therefore, if sdMAF were present, this sex-combined HWE test can be misleading. Instead of the Pearson’s 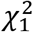 test, testing for model fit has also been proposed ^60^.

Regardless of the specific HWE test used, screening out variants with HWD is questionable for the X chromosome for two other reasons. First, it has been long (but not well) known that on the X chromosome, it takes several generations to achieve HWE, in contrast to a single generation for the autosomes, under the same set of assumptions such as random mating^22^. Second, for the autosomes, recent works^61–63^ have shown that association power can be improved by leveraging the *difference* in HWD between cases and controls, while remaining robust to HWD caused by genotyping errors, but this has yet to be explored for the X chromosome.

### X-inactivation uncertainty and association results interpretation

Until very recently, the statistical genetics community believed that X-inactivation was the main analytical challenge to achieving X chromosome-inclusive GWAS^1,11^. Therefore, most of the association methods developed so far have focused on X-inactivation^64–69^. As the true model can be no, random or skewed X-inactivation, the proposed methods include using minimum-p^64^, and model selection^66^, and Bayesian model averaging^69^.

However, it has been shown, both theoretically and empirically, that GWAS data alone cannot identify the true underlying X-inactivation model, due to the statistical confounding (e.g. skewed X-inactivation is confounded with non-additive genetic effect)^9^. While these observations helped to develop a new association test that is robust to X-inactivation uncertainty^9^, both SNP and sex effect estimates are biased if the model assumptions were incorrect^70^. As SNP effect estimates are the bases for constructing PGS or polygenic risk scores (PRS), future research should consider how to correct for the biases when X chromosomal variants are included in PRS.

### Heterogeneous reporting of summary statistics and the X chromosome results

There is a large variation in reporting standards of genome-wide summary statistics^71^. For example, among the 327 summary statistics files analyzed, there were 127 unique formats. The authors then developed MungeSumstats, a Bioconductor package to standardize and perform quality control of GWAS summary statistics. Interestingly, among the total of 31 checks listed in their Table 1, #27 checks “for SNPs on chromosome X, Y and mitochondrial SNPs, [and] if any are found these are removed,” even though an option of retaining them was provided.

We further examined the reporting standard in the original publications of the X chromosomal signals documented in the NHGRI-EBI GWAS catalog^5,6^ (downloaded on 2020-03-08 and initially reported in^72^) using the genome-wide significance level of p-value < 5 × 10^−8^. Out of the 3,869 studies available at that time (male-only studies excluded), 195 reported a total of 253 genome-wide significant loci on the X chromosome. To streamline the analysis, we selected only one SNP from each associated region by retaining the SNP with the smallest association p-value (Supplementary Data S2).

We then extracted information on the analyses performed from the original publications; in total there are 36 columns in Supplementary Data S2. These details are crucial to the analysis and reporting of the X chromosome, but are largely irrelevant to the autosomes. They include, for example, whether i) the analysis was sex-stratified (70% did sex-combined analysis), ii) for sex-combined analysis, sex was included as a covariate (57% did not), and iii) the genotype coding was documented (75% did not, presumably used the default X-inactivation assumption), because if X-inactivation was assumed, males are typically coded 0 and 2 respectively for the two hemizygous genotypes. These considerations were not included in the guidelines recommended by^73^. Not surprisingly, there was much heterogeneity in both the analysis and reporting among the 195 studies we examined. We suggest sex chromosome-aware research guidelines to be developed by the community.

### The Y chromosome

Non-recombing Y chromosome haplotypes have a long history in population genetics and genealogy^74^, since these haplotypes can be determined without ambiguity, making it the patrilineal equivalent to Mitochondrial haplogroups^75^. However, the Y chromosome has long been a thorn in the side of human geneticists: More than half of the Y chromosome is absent from GRCh38^76^. Two recent papers used combinations of multiple long-read next-generation sequencing technologies to generate much more complete sequence of the Y chromosome, and they also described a high degree of heterogeneity in chromosome length and content between individuals^76,77^.

The Telomere-to-Telomere consortium has reported the sequence of an approximately 62 Mb long human Y chromosome^76^, which includes >30Mb that were missing from the reference sequence. Human geneticists can often be criticized for exaggeration, claiming that their phenotype or gene of interest has extensive complexity, but the recent analysis of 43 diverse Y chromosomes takes the crown. For example, some Y chromosomes are only 45 Mb, while others are as long as 85 Mb, due in part to large duplications and inversions.

It has been shown that standard sequencing alignment methods may be problematic for females, without masking the Y chromosome from the reference genome^78^. For example, more variants were called after masking the Y chromosome in females, particularly in PAR regions. Similarly, for variant calling in PAR regions in males, it was recommended to provide only one PAR reference sequence from the two sex chromosomes (i.e. either the X or Y chromosome). Prior to variant calling, the authors recommended using read depths for the X and Y chromosomes, relative to the autosomes, to determine the sex chromosome composition of a sample, similar to that proposed for GWAS arrays^79^.

Additionally, in the past few years, age-dependent clonal loss of the Y chromosome has been reported in leukocytes^80,81^. This phenomenon may further affect data quality and analysis of PAR and Y chromosomal variants.

### Clinical Implications

The exclusion of sex chromosomes from analysis and reporting also has significant clinical implications. Chief among these is failure to identify disease-associated SNPs or regions important in pathophysiology, prevention, diagnosis, or treatment. While common GWAS-identified SNPs tend to have small estimated effect sizes, this is not necessarily true for SNPs affecting drug responses, which have not generally been subjected to strong selective pressures.

Current pharmacogenetic guidelines such as those of the Clinical Pharmacogenetics Implementation Consortium^82^ do not include genes on the sex chromosomes (quite possibly due to exclusion of these genes from analyses); were such variants to be identified, guidelines for screening or drug dosing might need to be modified based on a patient’s biologic sex. Similarly, adequate identification and inclusion of sex chromosome variants in polygenic risk scores might mandate stratification of these predictions by sex. It will be difficult, if not impossible, to assess these sex-stratified risks accurately until the dearth of analyses of sex chromosomes in clinically important traits is rectified.

## Recommendations and Future Directions

After 15 years, several authors have observed that GWAS are “realizing the promise”^83^, with “no signs of slowing down”^84^. Interestingly, sex chromosomes were not discussed in the 5-^85^, 10-^4^ and 15-^83,84^ reviews of GWAS. Our work here revealed that, for example, sex chromosomes are omitted from ~75% of the GWAS in 2021, which is likely the major cause of the paucity of signals on the sex chromosomes. Given these observations, to achieve sex chromosome-inclusive research, we make several recommendations and discuss related future research directions.

First, the existing bioinformatic and sequencing pipelines need to be revised for the sex chromosomes, from variant calling^78^ to imputation, so that the downstream analyses improve the integrity and robustness of sex chromosome analyses and provide greater confidence in conclusions drawn from them.

Second, quality control procedures need to follow previously recommended sex-stratified approaches^1,11^. Additionally, sex difference in MAF^52,86,87^ needs to be examined, but whether attributing significant sdMAF to genotyping errors (then screening out such variants) warrants future research. This is because sdMAF could also be a result of sex-specific linkage, particularly at the PAR-NPR boundaries^43^.

Third, the distinction is important between association testing and effect size estimation^9,70^. In particular, due to X-inactivation uncertainty, genetic effects cannot be reliably estimated, impacting future research on X chromosome-inclusive PRS-based risk prediction. As it is intractable to obtain genetically meaningful effect size estimates from sex-combined analysis based on GWAS data alone, it is thus reasonable to construct sex-specific PRS^88^, conceptually analogous to population-specific PRS^89^.

Fourth, obtaining and then incorporating SNP/gene/tissue/individual-specific X-inactivation could improve association methods. To this end, recent advances in long-read next generation sequencing technology, enabling phased allele-specific methylation, could be useful^90^.

Additionally, gene expression data such as the GTEx resource can be utilized^10,91^.

Fifth, many other existing statistical genetics analyses require sex chromosome-aware development and implementation. These include, for example, rare variants^92,93^, meta-analysis^94^, LD-score regression^95^, pleiotropy^96^, and causal inference via Mendelian Randomization^97^. More work is also need to better understand trait heritability^98^ attributed to the sex chromosomes.

In summary, ten years after the seminal work by ^1^, the exclusion of the X and Y chromosomes from whole-genome analysis persists. Until the sex chromosomes are indeed included in a whole-genome study, instead of GWAS, we propose they be more properly referred to as “AWAS” for “autosome-wide scan”.

## Acknowledgements

The authors would like to thank Karl Broman, Sara Good, Anthony Herzig, Inke König, Michael Schatz, Bhooma Thiruvahindrapuram, Melissa Wilson, and Andreas Ziegler for valuable discussions. This research was funded by the Canadian Institutes of Health Research (CIHR, PJT-180460), a University of Toronto Data Sciences Institute (DSI) Catalyst Grant.

## Author Contributions

LS, ADP and TAM conceptualized the study.

LS, ADP supervised the study and drafted the manuscript.

ZW and TL performed the analyses and summarized the results.

ZW, TL and TAM reviewed and edited the manuscript.

## Web Resources

Significant SNPs reported in the NHGRI-EBI GWAS catalog: https://www.ebi.ac.uk/gwas/docs/file-downloads

Genome-wide summary statistics reported in the NHGRI-EBI GWAS catalog: https://www.ebi.ac.uk/gwas/downloads/summary-statistics

The PolyGenic Score (PGS) catalog: https://www.pgscatalog.org/downloads/

## Data and Code Availability

All data used are publicly available. The specific downloads of the time-stamped datasets and codes used for the different analyses are all available at https://github.com/Paterson-Sun-Lab/eXclusionarY/

## Figure Legends

**Figure S1.**
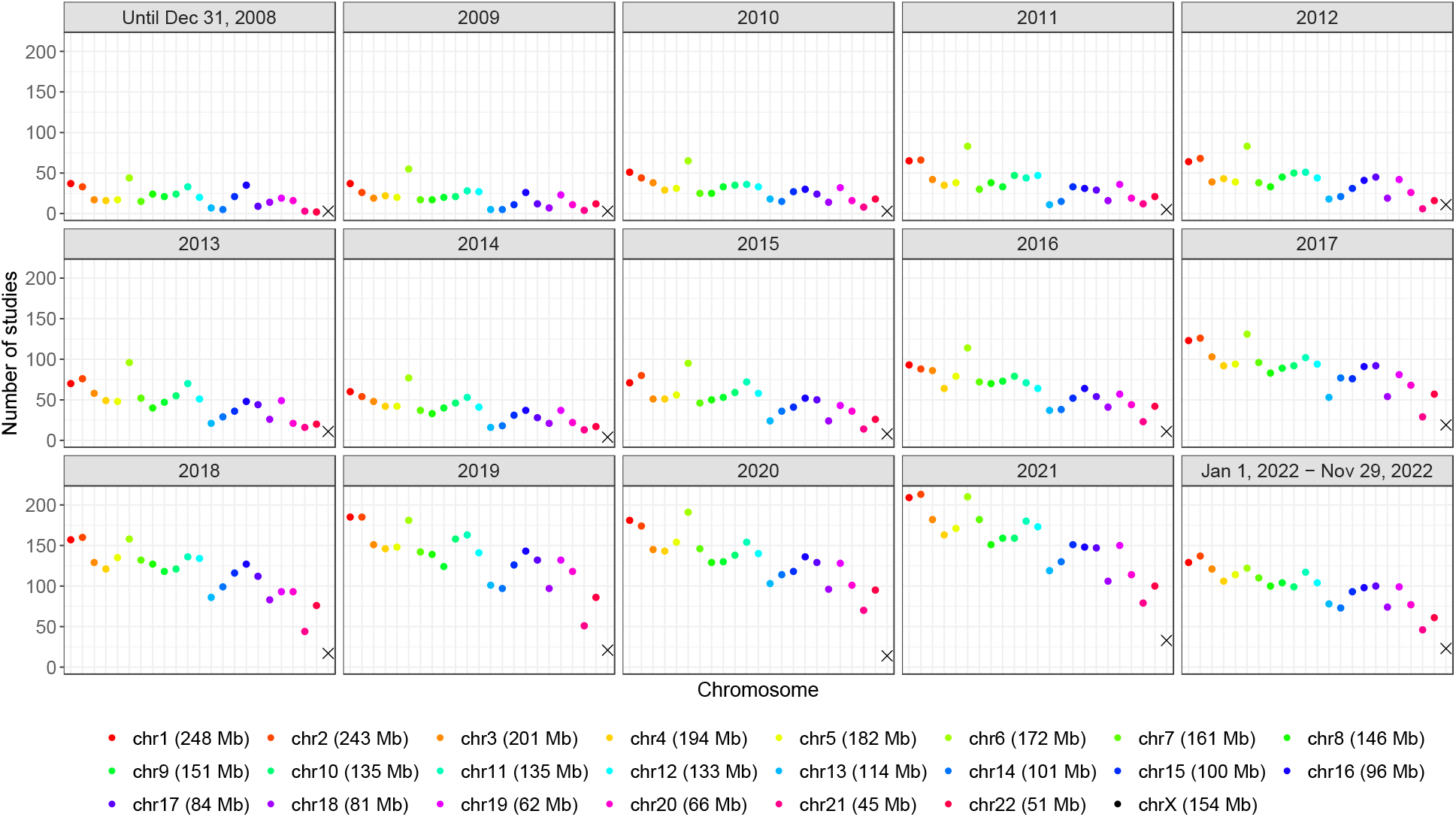
Total number of studies reporting at least one genome-wide significant finding (p-value < 5 × 10^-8^) over time, stratified by chromosome and year, from the NHGRI-EBI GWAS Catalog. Genetic associations were indexed by unique PubMed IDs. Studies reporting associations with multiple traits were only counted once.

**Figure S2.**
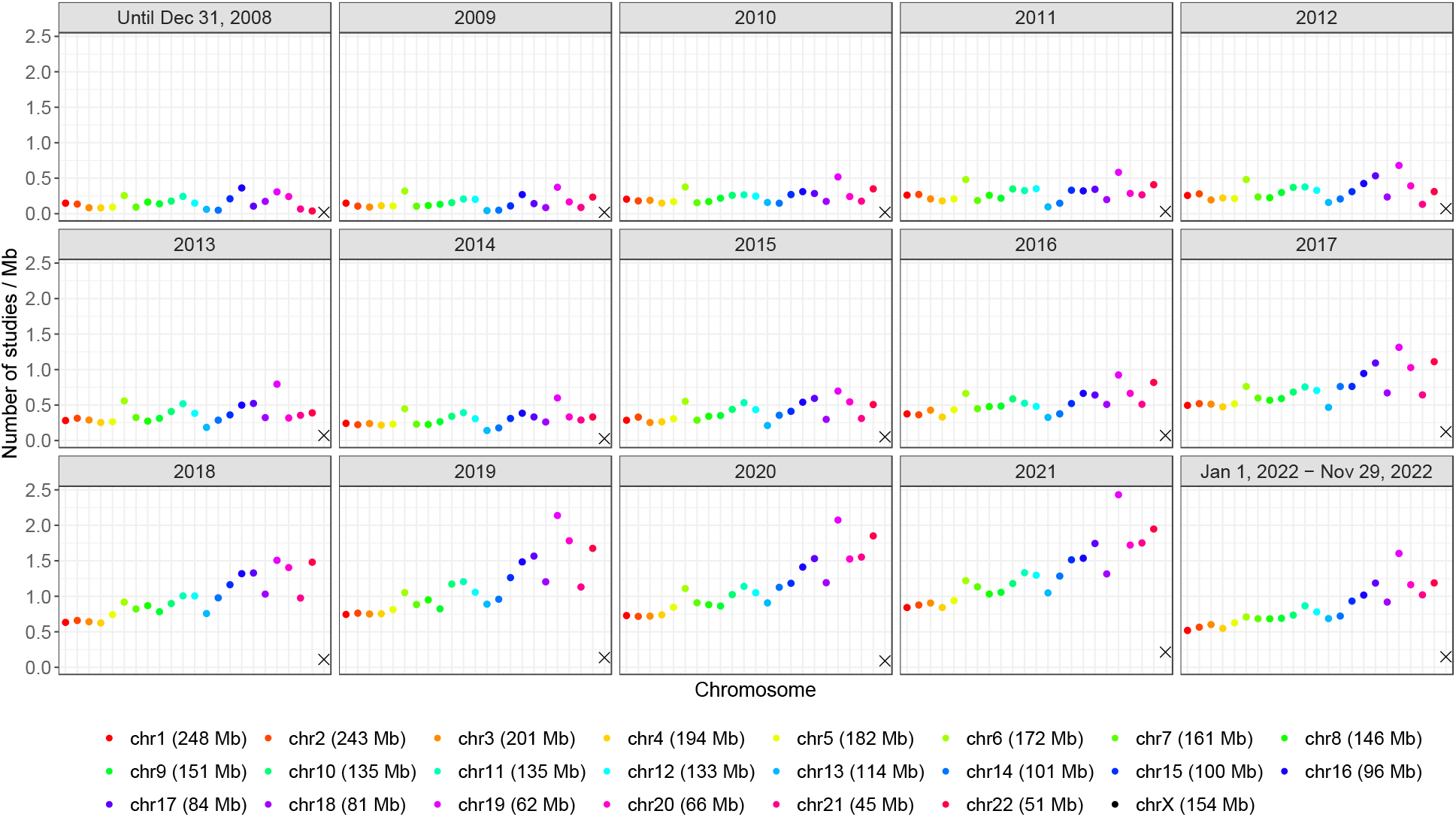
Average number of studies per Mb reporting at least one genome-wide significant finding (p-value < 5 × 10^-8^) over time, stratified by chromosome and year, from the NHGRI-EBI GWAS Catalog. Genetic associations were indexed by unique PubMed IDs. Studies reporting associations with multiple traits were only counted once.

